# Knockout of TSPO delays and reduces amyloid, Tau, astrocytosis and behavioral dysfunctions in Alzheimer’s disease

**DOI:** 10.1101/2022.03.26.485919

**Authors:** Kelly Ceyzériat, Léa Meyer, Farha Bouteldja, Stergios Tsartsalis, Quentin Amossé, Ryan J. Middleton, Guo-Jun Liu, Richard B. Banati, Thomas Zilli, Valentina Garibotto, Philippe Millet, Benjamin B. Tournier

**Affiliations:** Department of Psychiatry, University Hospitals of Geneva, Switzerland; Division of Nuclear medicine, Diagnostic Department, University Hospitals and CIBM Center for Biomedical Imaging, Faculty of Medicine, University of Geneva, Geneva, Switzerland; Division of Radiation Oncology, Department of Oncology, University Hospitals of Geneva, Switzerland; Australian Nuclear Science and Technology Organisation (ANSTO), Australia; Brain and Mind Centre, Medical Imaging Science, Faculty of Medicine and Health, The University of Sydney, Australia; Department of Psychiatry, University of Geneva, Switzerland

## Abstract

The 18kDa translocator protein (TSPO) is up-regulated in glial cells in neurodegenerative diseases. In Alzheimer’s disease (AD) animal models, TSPO is first overexpressed in astrocytes and then in microglia. However, the precise role of TSPO in the onset and progression of pathology and symptoms characteristic of the disease remains unknown. Here, we report that in the absence of TSPO in 3xTgAD mice the expected disease onset is significantly delayed and a reduction is seen in the hippocampal load of poorly and highly aggregated forms of Tau (−44% and −82%, respectively) and Aβ42 (−25% and −95%, respectively), at 9 months of age. In addition, the astrocyte reactivity was decreased in 3xTgAD.TSPO^−/−^ mice with a reduction in the morphologic complexity and the size of astrocytes in the dorso-dorsal hippocampus and the hilus. Functionally, the absence of TSPO ameliorated the cognitive consequences of adeno-associated virus-induced Tau over-expression in the hippocampus. This suggests that TSPO plays an important role in the active disease progression of AD. TSPO-inhibiting drugs thus merit further exploration as to their potential to reduce the rate of neurodegenerative disease progression.

## Introduction

Alzheimer’s disease (AD) is the most common form of age-related dementia and represents a very significant economic and social cost. It is characterized by the gradual accumulation of dysfunctional neurochemical markers including amyloid and Tau which lead to progressive loss of cognitive functions and brain atrophy^1^. The intraneuronal presence of both amyloid and Tau is well described and may be the basis of synaptic dysfunction and subsequent neuron death^1, 2, 3, 4, 5^. However, various therapies that have targeted amyloid or Tau have been unsuccessful^6^. Beyond the role of neurons in AD, the relative importance of glial cells is increasingly acknowledged. Astrocyte reactivity and microglial activation involve changes in their morphology and genetic signature^7, 8, 9^. These functional changes can be either protective or aggravating of the pathology depending on the area of the brain considered, the pathological stage and severity and the duration of their activation^10, 11^. A marker of their activation state during acute but also chronic neurodegenerative disease, is the measurement of the density of the 18kDa translocator protein (TSPO)^12^. TSPO, previously known as the peripheral benzodiazepine receptor, is present in the outer mitochondrial membrane^13^. It is accepted that TSPO is upregulated during pathology, in humans as well as in animal models including 3xTgAD mice^8^.

The cellular origin of TSPO is multiple, thus complicating the interpretation of variations in its density, although it has been shown that only glial cells over-express TSPO in AD^14, 15^. And, like other markers of reactivity, TSPO is first activated in astrocytes and then in microglia^16, 17, 18, 19^. Thus, even if TSPO appears to be a functional marker of glial activation, its specific involvement in AD remains largely unknown. The early increase of TSPO and the fact that its accumulation correlates with that of amyloid plaques, however, suggests a role in the development of the pathology. Previous studies tend to show that the absence of TSPO reduces the reactivity of glial cells and the inflammatory response^20, 21, 22^. Thus, to determine the function of TSPO in AD, the use of TSPO knockout mice can be very useful^23^. In this study, we investigated TSPO as a potential therapeutic target by examining its knockout in 3xTgAD mice. To measure the appearance and evolution of pathological markers, animals were studied at different ages: at 2 months, to assess the impact on the brain development; 4 months, simulating young adults and at the age of 9 months to mimic a very early stage of the pathology, when amyloid deposits are not yet observed in the hippocampus.

## Results

### TSPO colocalizes with amyloid deposits and influences amyloid and Tau accumulation

In the human AD brain, TSPO is associated with the presence of neuritic deposits formed by amyloid β (Aβ) and Tau (Fig. 1a). Its density is increased from the early stages of the pathology: hippocampus samples from Braak 6 subjects showed significantly higher levels than controls (p< 0.05) while Braak 4 subjects show intermediate levels (Fig. 1b). In addition, the accumulation of TSPO correlates with that of amyloid, suggesting that TSPO might be implicated in the mechanisms of the disease progression (Fig. 1c).

**Fig. 1.**
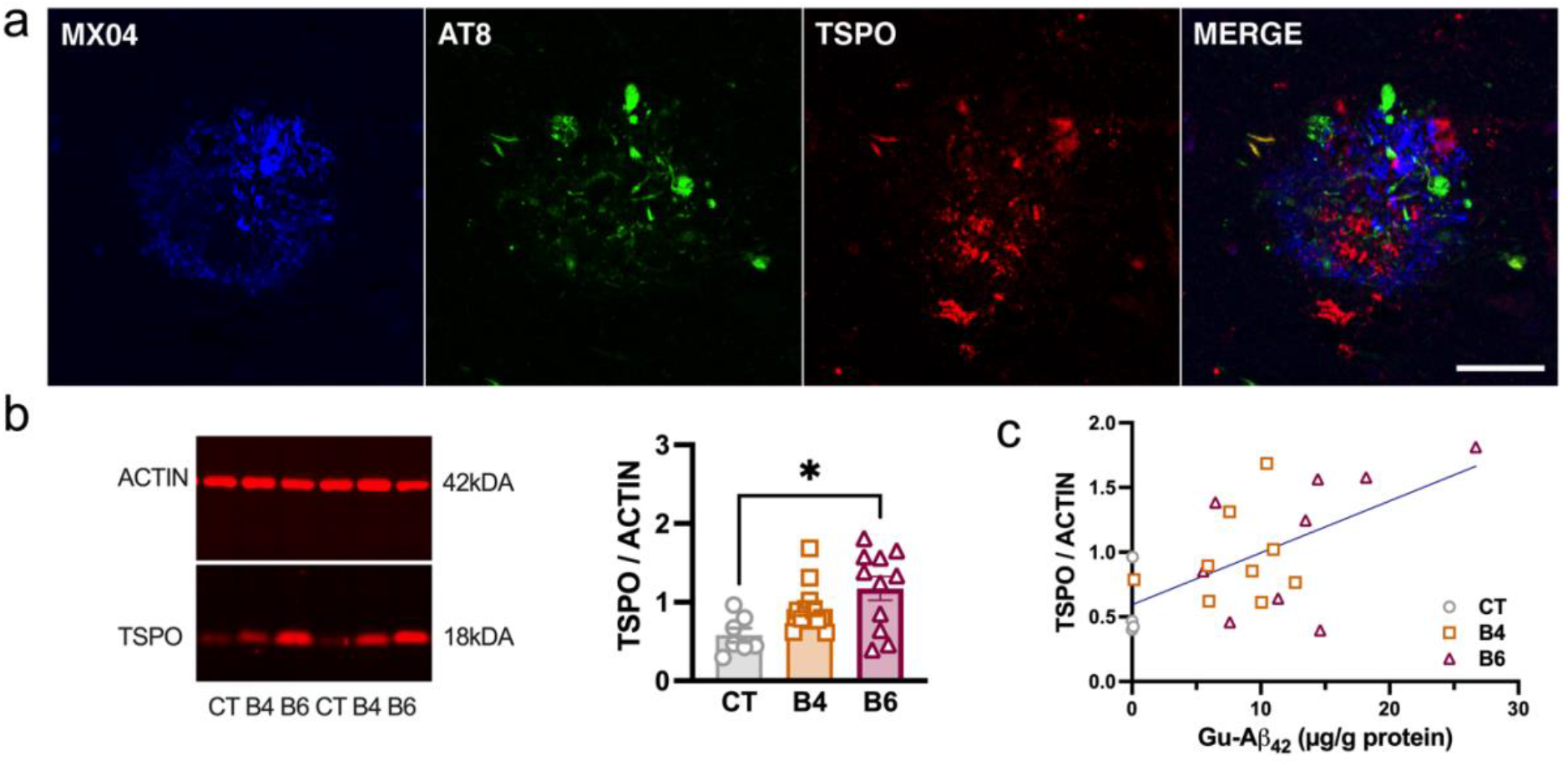
TSPO colocalizing with amyloid plaques and Tau is overexpressed in the hippocampus in human AD. **a**, Colocalization between Aβ(MXO4, blue), Tau (AT8, green), TSPO (red) in the human hippocampus. Scale bar: 50 μm. **b**, Quantification of TSPO in the hippocampus of control, Braak 4 (B4) and Braak 6 (B6) subjects. Data are presented as individual values and mean ±SEM (n=31) and analyzed by the Kruskal-Wallis test with the Dunn’s multiple comparisons *post hoc* test.

To determine whether TSPO plays a role in the etiopathology of AD, we knocked out TSPO in an AD mouse model by crossing TSPO^−/−^ with 3xTgAD mice, to create 3xTgAD.TSPO^−/−^ mice. The absence of TSPO is observed at the DNA and protein levels using gDNA PCR, western blotting (Fig. 2a,b) and *in situ* immunofluorescence staining (Fig. 2c).

**Fig. 2.**
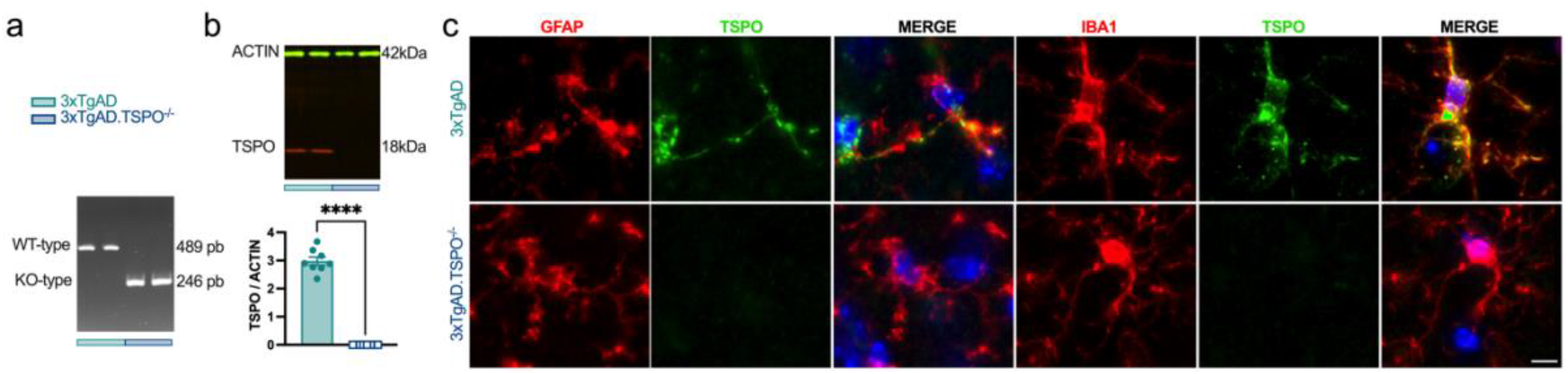
3xTgAD.TSPO^−/−^ mice lack TSPO in astrocytes and microglia, compared to 3xTgAD mice. **a**, gDNA PCR gel shows the discrimination of size of *Tspo* amplicon, validating the knock-out in 3xTgAD.TSPO^−/−^ mice. **b**, Measure of TSPO protein by western blot in the hippocampus confirmed the abolition of TSPO in 3xTgAD.TSPO^−/−^ mice (two-tailed unpaired t-test: *P* < 0.0001). **c**, Representative images of TSPO (green) staining in astrocytes (GFAP^+^) or in microglia (IBA1^+^). Scale bar: 20μm.

Immunological labeling of amyloid with 6E10 and A4G8 antibodies reveals, depending on the age of the animals, a difference between the genotypes. Indeed, hippocampal pyramidal neurons showed a greater accumulation of 6E10 in the absence of TSPO at 9 months of age (Fig. 3a,b). The same observation is made by measuring the intensity of the A4G8 staining with, in addition, a decrease at the age of 4 months (Fig. 3c). No extracellular Aβ deposit was observed in mice aged 2, 4 and 9 months, and granule neurons of the hippocampal dentate gyrus were also negative for 6E10 and A4G8. We then quantified the levels of amyloid in the parenchyma by by separating the proteins solubilized in triton (Tx) and guanidine (Gu) which represent the non or weakly aggregated forms and the more strongly aggregated forms, respectively. Data shows a decreased amyloid accumulation in 3xTgAD.TSPO^−/−^ mice (Fig. 3d). Specifically, at the age of 9 months a decrease in Tx-Aβ42 is observed (−26 ±3.7%, p <0.001) and Gu-Aβ42 is almost absent (−96 ±0.8%, p <0.0001). Unlike Aβ42, which is the most toxic form of the amyloid, the accumulation of Aβ40 is not significantly affected by the absence of TSPO. To analyze the reasons for this decrease, we measured the density of proteins involved in the production and degradation of amyloid. The levels of BACE1 (β-secretase), ApoE (ApolipoproteinE) and IDE (insulin-degrading enzyme) were quantified by western blot in the hippocampus and did not show differences between genotypes (Fig. 3e). In addition, APP mRNA levels and the soluble APPα/ soluble APPβ ratio were not different between the genotypes (Fig. 3f, g).

**Fig. 3.**
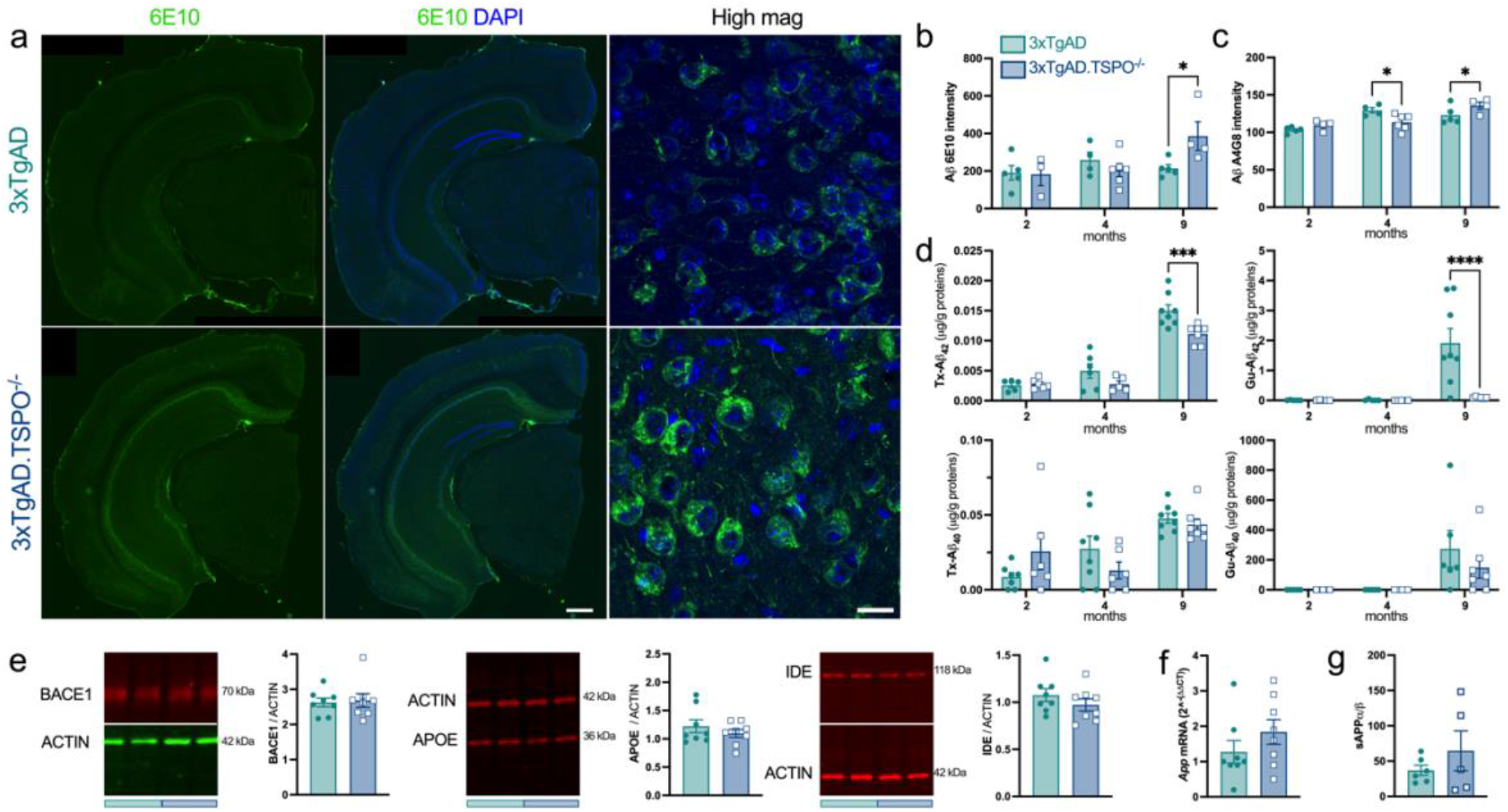
3xTgAD.TSPO^−/−^ mice exhibit reduced and delayed amyloid pathology. **a**, Representative example of 6E10 perinuclear immunoreactivity (ir-, green) and DAPI (blue) in 9-month-old 3xTgAD and 3xTgAD.TSPO^−/−^ mice. Scale bar: 500 μm and 20 μm in high magnification images. **b**, Quantification of 6E10-ir perinuclear labeling (two-way ANOVA: genotype x age, *F_2,22_* = 6.59, *P* = 0.0057; genotype, *F_1,22_* = 0.07, *P* > 0.05; age, *F_2,22_* = 14.92, *P* < 0.0001). **c**, Quantification of A4G8-ir intensity (two-way ANOVA: genotype x age, *F_2,21_* = 3.60, *P* = 0.045; genotype, *F_1,21_* = 1.00, *P* > 0.05; age, *F_2,21_* = 3.07, *P* = 0.067). **d**, Quantification of triton x100- (Tx) and guanidine- (Gu) soluble forms of Aβ40 and Aβ42 (two-way ANOVA, Tx-Aβ40: genotype x age, *F_2,37_* = 2.79, *P* > 0.05; genotype, *F_1,37_* = 0.006, *P* > 0.05; age, *F_2,37_* = 11.91, *P* = 0.0001; Tx-Aβ42: genotype x age, *F_2,31_* = 3.33, *P* = 0.049; genotype, *F_1,31_* = 8.66, *P* = 0.0061; age, *F_2,31_* = 109.5, *P* < 0.0001; Gu-Aβ40: genotype x age, *F_2,28_* = 0.67, *P* > 0.05; genotype, *F_1,28_* = 0.63, *P* > 0.05; age, *F_2,28_* = 7.64, *P* = 0.0023; Gu-Aβ42: genotype x age, *F_2,28_* = 7.13, *P* = 0.003; genotype, *F_1,28_* = 6.53, *P* = 0.016; age, *F_2,28_* = 8.63, *P* = 0.0012). **e**, Quantification of protein in the whole hippocampus. From left to right, representative immunoblot (showing two samples from 3xTgAD and two samples from 3xTgAD.TS PO^−/−^ mice) and protein quantification (normalized to ACTIN) for BACE1, APOE and IDE (two-tailed unpaired t-test: p>0.05). **f**, mRNA quantification of *App* (2^^^-^ΔΔCT^ method; two-tailed unpaired t-test: p>0.05). **g**, soluble APPα/ soluble APPβ ratio (two-tailed unpaired t-test: p>0.05).

We next examined if the absence of TSPO induces changes in the second AD hallmark, namely phosphorylated-Tau (Fig. 4a). Immunoreactivity for AT8 showed a delay in the appearance of AT8 positive neurons in the pyramidal cell layer (Fig. 4b). Indeed, at 4-month-old, 3xTgAD.TSPO^−/−^ mice are less prone to express Tau (p=0.009) and a trend persists at 9 months of age (p=0.09). In addition, Tx-Tau and Gu-Tau levels are also affected by the absence of TSPO (Fig. 4c). In detail, a decrease in the concentration of Tx-Tau forms is observed at 4 (−45 ±11%, p<0.05) and 9 (−44 ±8.9%, p<0.05) months. While the Gu-Tau forms are observed at 9 months of age in 3xTgAD mice, they are strongly reduced in 3xTgAD.TSPO^−/−^ (−82 ±3.4%, p<0.0001).

**Fig. 4.**
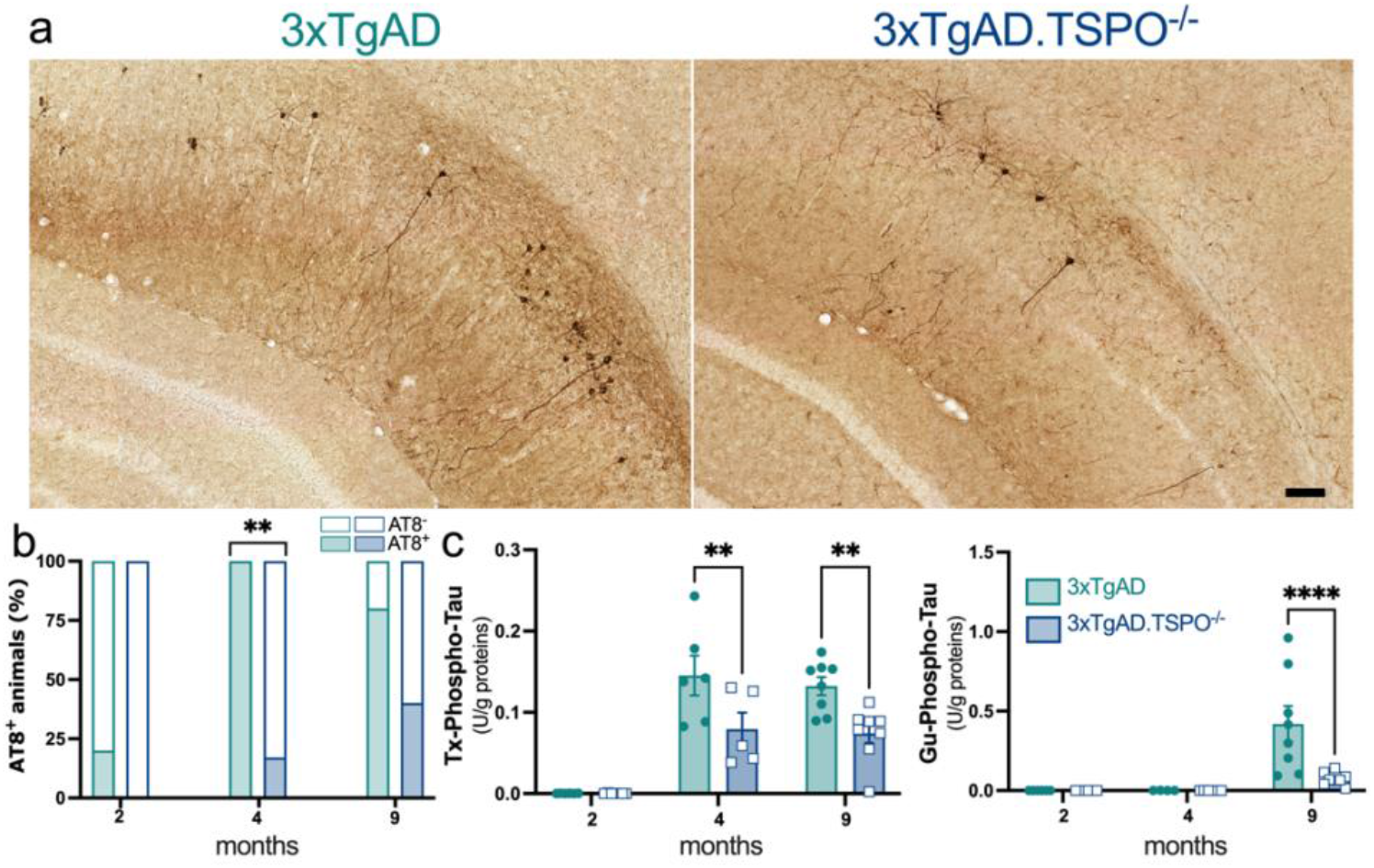
3xTgAD.TSPO^−/−^ mice exhibit reduced and delayed Tau pathology. **a**, Representative example of AT8-ir in 9-month-old 3xTgAD and 3xTgAD.TSPO^−/−^ mice. Scale bar: 100μm. **b**, Number of mice AT8 positive as function of the age (chi-squared, *P* < 0.01). **c**, Quantification of Triton- (Tx) and guanidine- (Gu) soluble forms of pT231 Tau (two-way ANOVA, Tx-Tau: genotype x age, *F_2,34_* = 3.47, *P* = 0.04; genotype, *F_1,34_* = 13.84, *P* = 0.0007; age, *F_2,34_* = 41.56, *P* < 0.0001; Gu-Tau: genotype x age, *F_2,32_* = 6.07, *P* = 0.006; genotype, *F_1,32_* = 5.29, *P* = 0.028; age, *F_2,32_* = 12.47, *P* < 0.0001).

### TSPO knockout reduces astrocyte reactivity in the hippocampus: a subdivision-dependent effect

Activation of glial cells is a widely studied phenomenon in human AD and the 3xTgAD model^24, 25^. To determine whether TSPO knockout modifies the functioning of astrocytes, we firstly measured CLUSTERIN, secreted by astrocytes and known to promote formation of extracellular Aβ deposits^26, 27^. At mRNA level, *Clu* is not affected but at the protein level, CLUSTERIN is significantly reduced in 3xTgAD.TSPO^−/−^ mice (−9 ±1.2 %, p<0.05, Fig. 5a), suggesting a reduction in the pro-inflammatory state of astrocytes^28^. In contrast, the absence of TSPO did not modify neither the expression nor the density of GFAP in the whole hippocampus. To go further in the investigation of astrocyte reactivity, an analysis was separately conducted in the subiculum and dorsal and ventral sub-divisions of the hippocampus, as previous studies have shown that the reactivity of astrocytes, the presence of AD markers (Aβ, Tau) and the expression of TSPO are heterogeneous depending on the location in the hippocampus^29, 30, 31^. Representative images of GFAP immunoreactivity and quantification in hippocampal areas are displayed in Fig. 5b,c. In the subiculum, the area occupied by astrocytes was clearly reduced in 3xTgAD.TSPO^−/−^ mice at 4 and 9 months of age (−65 ±1.6% and −67 ±5.6%, respectively). At 9 months, the same observation was made in the dorsal hippocampus (−50 ±7.5%). Astrocytes in the ventral hippocampus were not affected by the absence of TSPO. To validate that these effects are specific for 3xTgAD.TSPO^−/−^ mice, a comparative analysis was carried in control and TSPO^−/−^ mice. In this case, no WT vs TSPO^−/−^ difference was observed (Supp data Fig. 1).

**Fig. 5.**
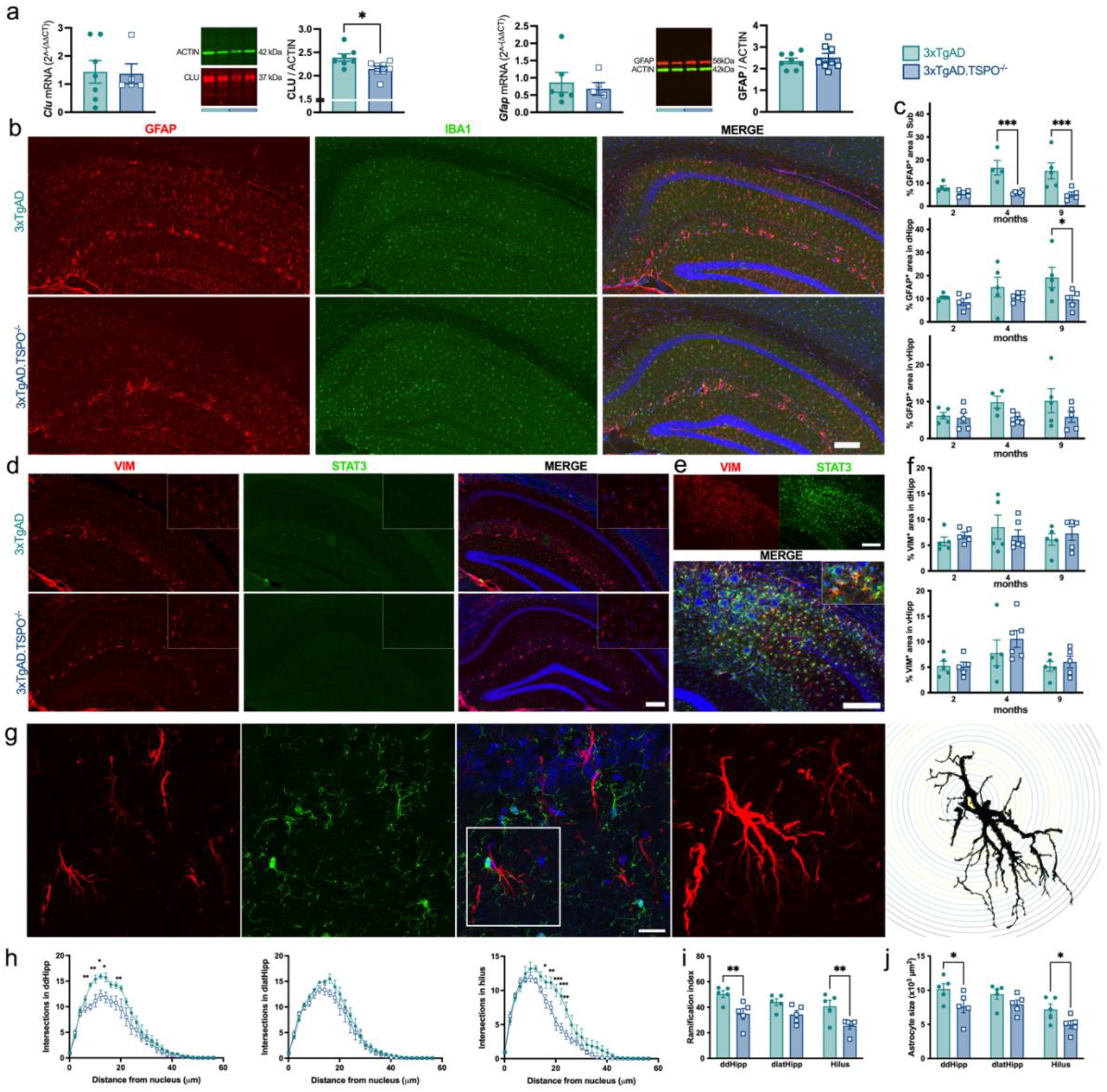
3xTgAD.TSPO^−/−^ mice exhibit reduced astrocyte reactivity. **a**, Quantification of mRNA and protein in the whole hippocampus. From left to right, mRNA quantification (2^^^-^(ΔΔCT)^), representative immunoblot (showing two samples from 3xTgAD and two samples from 3xTgAD.TSPO^−/−^ mice) and protein quantification (normalized to ACTIN) for GFAP, VIMENTIN (VIM) and CLUSTERIN (CLU) (two-tailed unpaired t-test: p>0.05). **b**, Representative example of GFAP (red), IBA1 (green) immunoreactivity and DAPI (blue in merge images) in 9-month-old 3xTgAD and 3xTgAD.TSPO^−/−^ mice. Scale bar: 200 μm. **c**, Quantification of % of positive GFAP-ir area in the subiculum (Sub), dorsal hippocampus (dHipp) and ventral hippocampus (vHipp), and intensity of the GFAP-ir staining the subiculum (%GFAP^+^, two-way ANOVA, Subiculum: genotype x age, *F_2,24_* = 3.16, *P* = 0.06; genotype, *F_1,24_* = 26.35, *P* < 0.0001; age, *F_2,24_* = 3.03, *P* = 0.06; dHipp: genotype x age, *F_2,25_* = 1.06, *P* > 0.05; genotype, *F_1,25_* = 6.19, *P* = 0.019; age, *F_2,25_* = 1.75, *P* > 0.05; vHipp: genotype x age, *F_2,24_* = 0.83, *P* > 0.05; genotype, *F_1,24_* = 5.07, *P* = 0.034; age, *F_2,24_* = 0.86, *P* > 0.05; GFAP^+^ intensity, two-way ANOVA, Subiculum: genotype x age, *F_2,22_* = 3.00, *P* = 0.07; genotype, *F_1,22_* = 34.95, *P* < 0.0001; age, *F_2,22_* = 3.32, *P* = 0.055). **d**, Representative example of VIM (red), STAT3 (green) immunoreactivity and DAPI (blue in merge images) in 9-month-old 3xTgAD and 3xTgAD.TSPO^−/−^ mice. Scale bar: 200 μm (inserts display high magnification). **e**, Positive control for VIM (left), STAT3 (right) immunoreactivity and merge images with DAPI (20-month-old 3xTgAD mouse). Scale bar: 200 μm. **f**, Quantification of % of positive VIM-ir area in dHipp and vHipp (%VIM^+^, two-way ANOVA, dHipp: genotype x age, *F_2,25_* = 0.78, *P* > 0.05; genotype, *F_1,25_* = 0.04, *P* > 0.05; age, *F_2,25_* = 0.56, *P* > 0.05; vHipp: genotype x age, *F_2,25_* = 0.49, *P* > 0.05; genotype, *F_1,25_* = 0.92, *P* > 0.05; age, *F_2,25_* = 4.36, *P* = 0.024). **g**, Representative example of the signal extraction for Sholl analysis (GFAP, red; IBA1, green; DAPI, blue). Image on the right shows a high magnification of the insert. Scale bar: 20 μm. **h**, Quantification of the number of intersections as function of the distance from soma in dorso-dorsal (ddHipp), dorso-lateral (dlatHipp) hippocampus and hilus (two-way ANOVA, ddHipp: genotype x distance, *F_28,224_* = 3.02, *P* < 0.0001; genotype, *F_1,8_* = 5.34, *P* = 0.049; distance, *F_28,224_* = 129.7, *P* < 0.0001; dlatHipp: genotype x distance, *F_28,224_* = 0.88, *P* = 0.63; genotype, *F_1,8_* = 1.1, *P* > 0.05; distance, *F_28,224_* = 132.9, *P* < 0.0001; hilus: genotype x distance, *F_28,224_* = 4.15, *P* < 0.0001; genotype, *F_1,8_* = 5.96, *P* = 0.04; distance, *F_28,224_* = 151.9, *P* < 0.0001). **i**, Total number of ramifications (two-way ANOVA: genotype x area, *F_2,24_* = 0.46, *P* > 0.05; genotype, *F_1,24_* = 22.87, *P* < 0.001; area, *F_2,24_* = 3.80, *P* = 0.036). **j**, Size of the soma of astrocytes (two-way ANOVA: genotype x area, *F_2,24_*= 0.26, *P* > 0.05; genotype, *F_1,24_* = 11.4, *P* = 0.0025; area, *F_2,24_* = 8.71, *P* = 0.0014).

In addition to GFAP, two other markers of reactive astrocytes were studied, namely VIMENTIN and the canonical inducer of astrogliosis STAT3 (signal transducer and activator of transcription 3). At 9-month-old, 3xTgAD and 3xTgAD.TSPO^−/−^ mice are dimly positive for VIMENTIN and negative for STAT3 (Fig. 5d) in comparison with the clear STAT3 and VIMENTIN staining in positive old 3xTgAD control animals (Fig. 5e), indicating the very early stage of our study^32^. In addition, the VIMENTIN positive area did not differ between groups in the dorsal and the ventral hippocampus (Fig. 5f, the VIMENTIN signal in the subiculum was too low to be quantified).

To reveal astrocyte changes more thoroughly, we examined their morphology using sholl analysis (Fig. 5g). The complexity of the astrocytes (i.e. the number of branches according to the distance from the soma) was reduced in 3xTgAD.TSPO^−/−^ mice at the level of the dorso-dorsal hippocampus and the hilus but not at the level of the dorso-lateral hippocampus (Fig. 5h). The total branching number and the size of astrocyte soma were also significantly reduced in the dorso-dorsal hippocampus and hilus of 3xTgAD.TSPO^−/−^ mice (Fig. 5i,j), confirming a reduced activation of astrocytes.

### Absence of TSPO does not affect microglial cell density and morphology

To determine if microglia was also affected by the genetic knockout of TSPO, region of interest and cell level quantifications were performed using the IBA1 staining. The representative example of IBA1 staining in the Fig. 5b suggests the absence of genetic effect that was confirmed by the quantification in subiculum, dorsal and ventral hippocampus (Fig. 6a). Sholl analysis did not reveal any difference between genotype, whatever the subregion of the hippocampus, indicating that the microglial complexity (Fig. 6b,c) and size (Fig. 6d) was not impacted by TSPO knockout, at least at 9 months of age in 3xTgAD mice. The comparison of microglia between TSPO^−/−^ and WT mice did not reveal any difference (Supp data Fig. 1).

**Fig. 6.**
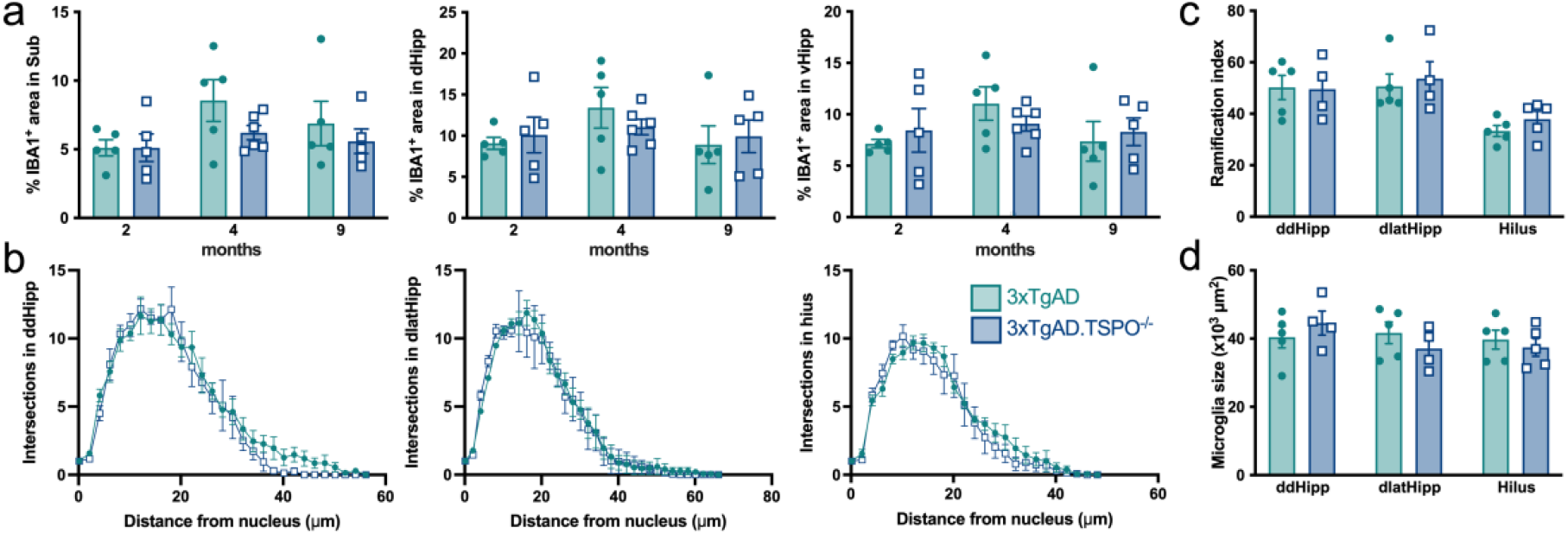
3xTgAD.TSPO^−/−^ mice did not show microglial activity changes. **a**, Quantification of % of positive IBA1-ir area in the subiculum (Sub), dorsal hippocampus (dHipp) and ventral hippocampus (vHipp) (%IBA1^+^, two-way ANOVA, Subiculum: genotype x age, *F_2,25_* = 0.60, *P* > 0.05; genotype, *F_1,25_* = 1.89, *P* > 0.05; age, *F_2,25_* = 2.26, *P* > 0.05; dHipp: genotype x age, *F_2,25_* = 0.57, *P* > 0.05; genotype, *F_1,25_* = 0.004, *P* > 0.05; age, *F_2,25_* = 1.51, *P* > 0.05; vHipp: genotype x age, *F_2,25_* = 0.76, *P* > 0.05; genotype, *F_1,25_* = 0.005, *P* > 0.05; age, *F_2,25_* = 1.65, *P* > 0.05). **b**, Quantification of the number of intersections as function of the distance from soma in dorso-dorsal (ddHipp), dorsolateral (dlatHipp) hippocampus and hilus using the Sholl analysis (two-way ANOVA, ddHipp: genotype x distance, *F_28,196_* = 0.64, *P* > 0.05; genotype, *F_1,7_* = 0.44, *P* > 0.05; distance, *F_28,196_* = 78.9, *P* < 0.0001; dlatHipp: genotype x distance, *F_33,231_* = 0.37, *P* > 0.05; genotype, *F_1,7_* = 0.02, *P* > 0.05; distance, *F_33,231_* = 86.8, *P* < 0.0001; hilus: genotype x distance, *F_24,192_* = 0.74, *P* > 0.05; genotype, *F_1,8_* = 0.24, *P* > 0.63; distance, *F_24,192_* = 72.9, *P* < 0.0001). **c**, Total number of ramifications (two-way ANOVA: genotype x area, *F_2,22_* = 0.18, *P* > 0.05; genotype, *F_1,22_* = 0.39, *P* > 0.05; area, *F_2,22_* = 8.06, *P* = 0.0024). **j**, Size of the soma of microglia (two-way ANOVA: genotype x area, *F_2,22_* = 1.06, *P* > 0.05; genotype, *F_1,22_* = 0.14, *P* > 0.05; area, *F_2,22_* = 0.91, *P* > 0.05).

### TSPO alters anxiety levels in the 3xTgAD mice

Among the early behavioral signs of AD, the presence of increased anxiety levels is widely reported, and even before the onset of cognitive impairment, in both human and 3xTgAD mice^33, 34^. As the mice used here at 9 months of age represent a very early stage of the disease, evaluation of anxiety and spatial working memory was performed.

The absence of TSPO leads to an increase in the time spent exploring open arms and performing head-dipping (Fig. 7a-c), demonstrating a decrease anxiety in 3xTgAD.TSPO^−/−^ mice. No difference was observed in terms of spatial working memory (Fig. 7d,e). Importantly, TSPO^−/−^ mice did not differ from WT mice (Supp data Figure 1).

**Fig. 7.**
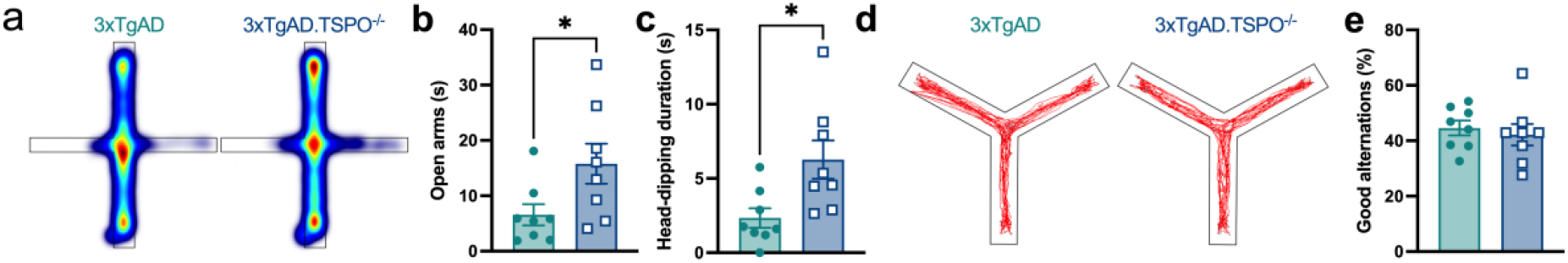
3xTgAD.TSPO^−/−^ mice show reduced levels of anxiety. **a**, Average heatmap of the exploring time (i.e. when the animal is moving) in the elevated plus maze (EPM). **b**, Time past in open arms in the EPM (two-tailed unpaired t-test: *P* = 0.0408). **c**, Head-dipping duration in the EPM (two-tailed unpaired t-test: *P* = 0.0167). **d**, Representative example of a mouse tracing during the 5 min of exploration of the Y-maze. **e**, % of good alternations in the Y-maze (two-tailed unpaired t-test: *P* > 0.05).

### TSPO knockout protects against Tau build-up and Tau-induced cognitive deficits

As cognitive decline in AD patients is better correlated to Tau pathology than Aβ, to evaluate if TSPO knock out directly affects Tau-related neuropathology and cognitive deficits, independently of amyloid pathology, we performed a supplemental experiment. WT and TSPO^−/−^ animals received an intra-hippocampal injection of adeno-associated virus (AAV) allowing the overexpression of Tau or eGFP (as a control). At 1- and 7-months post-Tau/eGFP induction, animals were tested for anxiety and spatial working memory to assess early and late effects. A reduction of anxiety (exploration time of open arms, duration of head-dipping) was induced by Tau, from 1 month after injection, but this effect was limited to TSPO^−/−^ animals (Fig. 8a,b). In addition, AAV-Tau induced a late decrease in the % alternation in the Y-maze test in WT mice (−36.74 ±8.06%, p<0.01), suggesting a deficit in spatial working memory (Fig. 8c). Importantly, TSPO knockout protects from the deleterious effect of Tau (−12.56 ±8.42%. p>0.05). *Postmortem* analysis validated the presence of Tau using HT7-immunoreactivity (Fig. 8d) and demonstrated the reduction in pT321 Tau-immunoreactivity in the hippocampus of AAV-Tau treated TSPO^−/−^ mice, as compared to WT (Fig. 8e).

**Fig. 8.**
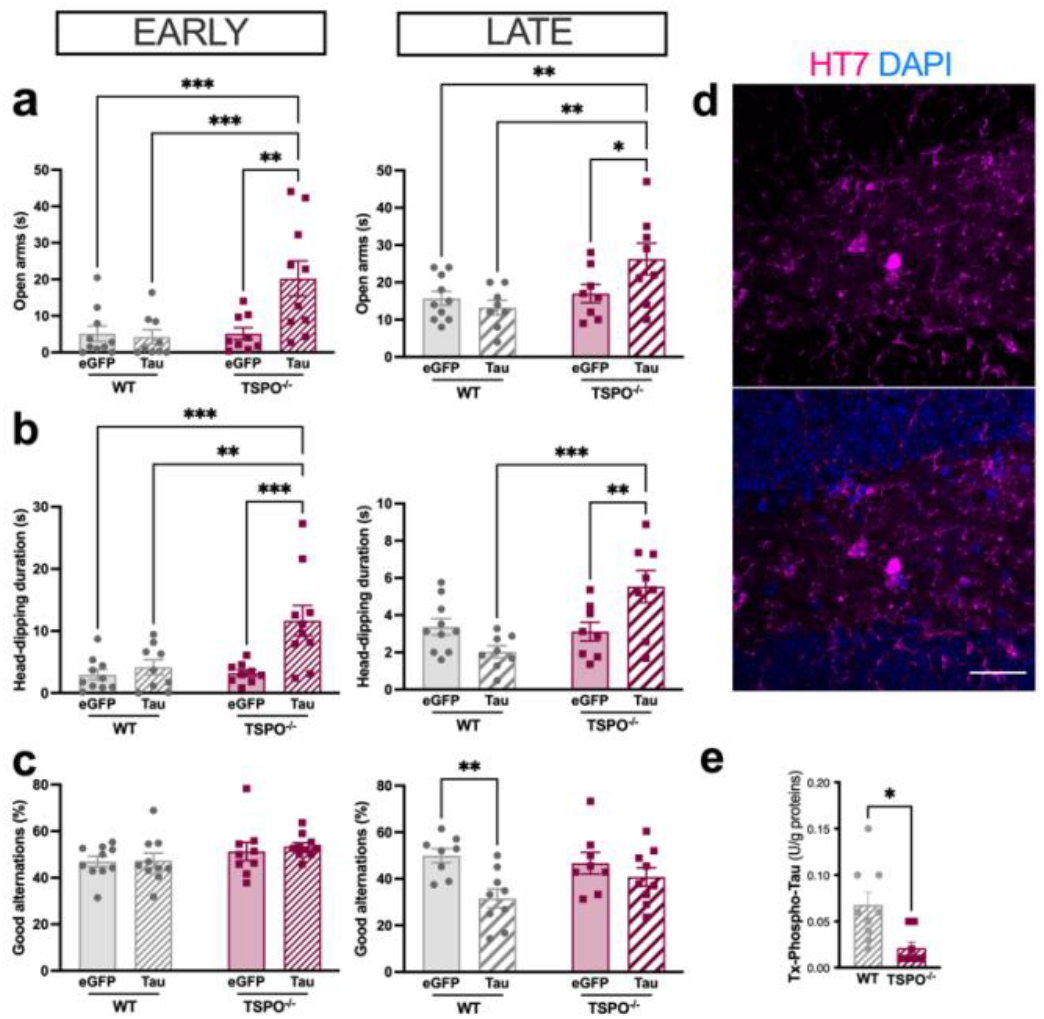
TSPO^−/−^ mice show reduction in Tau-induced phospho-Tau and cognitive deficit. Early (1 month) and late (6 months) behavioral effects induced by the intra-hippocampal injection of AAV encoding Tau of eGFP (control) gene. **a**, Time past in open arms in the EPM at 1 month (left: two-way ANOVA, genotype x AAV, *F_1,34_* = 6.93, *P* = 0.012; genotype, *F_1,34_* = 6.96, *P* = 0.012; AAV, *F_1,34_* = 5.47, *P* = 0.025) and 6 months (right: two-way ANOVA, genotype x AAV, *F_1,30_* = 4.61, *P* = 0.04; genotype, *F_1,30_* = 6.88, *P* = 0.013; AAV, *F_1,30_* = 1.56, *P* > 0.05) post Tau induction. **b**, Head-dipping duration in the EPM at 1 month (left: two-way ANOVA, genotype x AAV, *F_1,34_* = 5.9, *P* = 0.02; genotype, *F_1,34_* = 6.79, *P* = 0.013; AAV, *F_1,34_* = 10.4, *P* = 0.008) and 6 months (right: two-way ANOVA, genotype x AAV, *F_1,30_* = 11.26, *P* = 0.002; genotype, *F_1,30_* = 8.48, *P* = 0.007; AAV, *F_1,30_* = 0.93, *P* > 0.05) post Tau induction. **c**, % of good alternation in the Y-maze at 1 month (left: two-way ANOVA, genotype x AAV, *F_1,35_* = 0.09, *P* > 0.05; genotype, *F_1,35_* = 3.4, *P* > 0.05; AAV, *F_1,35_* = 0.2, *P* > 0.05) and 6 months (two-way ANOVA, genotype x AAV, *F_1,30_* = 2.45, *P* > 0.05; genotype, *F_1,30_* = 0.56, *P* > 0.05; AAV, *F_1,30_* = 9.26, *P* = 0.005; two-tailed unpaired t-test: AAV-Tau in WT: *P* = 0.0028; two-tailed unpaired t-test AAV-Tau in TSPO^−/−^: *P* = 0.34) post Tau induction. **d**, Representative example of Tau immunofluorescence at 6 months post Tau induction (top: Tau HT7 immunoreactivity; bottom: merge image with DAPI staining). Scale bar: 30 μm. **e**, Quantification of pT231 Tau (two-tailed unpaired t-test: *P* = 0.0104).

## Discussion

The use of TSPO as an *in vivo* marker of neuroinflammation in AD is widely reported in animal models and in humans^8, 12^, but its role in the etiopathology remains largely unknown. We show here that TSPO could represent a particularly interesting target for the treatment of AD. Indeed, its absence leads to a decrease in neuropathological markers (Aβ et Tau), to a decrease in astrocyte reactivity, an improvement in one of the early symptoms of AD (i.e. anxiety) and a decrease in Tau-induced cognitive dysfunctions.

At 9 months of age, a stage when no extracellular Aβ-plaque deposition was observed, there was an increase in the intracellular staining of Aβ as revealed by 6E10- and 4G8-immunoreactivity (-ir). In contrast, we reported a decrease in secreted forms of Aβ42 (poorly and highly aggregated). These effects could be explained by an alteration in the ratio of the metabolism of APP between the non-amyloid and the amyloid pathways. Indeed, APP can be cleaved by an alpha-secretase which generates a soluble amino-terminal fragment alpha (sAPPα) that prevents the generation of Aβ. APP can also be cleaved by a β-secretase allowing the generation of the Aβ40 and Aβ42 fragments. However, 6E10-ir is, at least in part, a marker of sAPPα while Aβ40-ir and Aβ42-ir represent the pathological forms of APP catabolism. Our data showing increase in 6E10-ir and decrease in Aβ42-ir seem to indicate an improvement in the imbalances in the processing of APP towards the non-amyloidogenic pathway. In addition, the direct consequence of the accumulation of sAPPα being a limitation of the amyloidogenic maturation of APP^35^, increase of sAPPα in 3xTgAD.TSPO^−/−^ mice may therefore amplify the reduction of secreted Aβ. However, the density of BACE1 remains unchanged, indicating the maintenance at least in part of the Aβ synthesis pathway, unless BACE1 activity is reduced. To assess if BACE1 activity is modified, the sAPPα/ sAPPβ was estimated but did not show a significant effect. Unlike impaired amyloid production, the levels of Aβ degrading enzymes ApoE and IDE were found to be unaffected by TSPO knockout. However, other mechanisms could come into play, as for example, the phagocytosis capacities of glial cells, the first key element in the degradation of Aβ.

Beyond Aβ, TSPO knockout also leads to a drastic reduction in Tau pathology. The accumulation of Tau in humans seems to be more related to the onset and course of cognitive decline^36, 37^, but at the age of 9 months, 3xTgAD mice do not yet show impaired spatial working memory. In contrast, they show signs of anxiety, one of the markers of early neuropsychiatric symptoms of AD^38, 39^. Consistent with the idea of an ameliorating effect of the absence of TSPO, mice show reduced anxiety levels. Moreover, to further analyze the impact of TSPO on Tau pathology, we show a functional relationship between these two actors. Indeed, the exogenous overproduction of Tau induces higher densities of phospho-Tau and a highest cognitive impairment in the presence of TSPO. This suggests that the absence of TSPO limits Tau phosphorylation which would in turn limit cognitive loss. In an observational study, the accumulation of TSPO was shown to be correlated with that of phospho-Tau in P301S mice^40^, which supports our results. This effect may be related to the reduction in astrocyte reactivity observed in 3xTgAD.TSPO^−/−^mice.

Indeed, at the early stages used herein, we observed an impact of the absence of TSPO on the astrocytes but not on the microglia. The absence of microglia changes in 3xTgAD.TSPO^−/−^ mice resembles to the unimpaired activation of microglia in TSPO^−/−^ mice in response to neuronal injury^23^ and could indicate the relative low function of TSPO in microglia. However, TSPO appears to be required for microglia-spine interactions in response to diazepam^20^, suggesting a role for microglial TSPO later in AD pathology. Considering astrocytes, the lack of difference at 2 months of age and the lack of differences between WT and TSPO^−/−^ mice seem to prove that the reduction in astrocyte reactivity is not innate but is the consequence of the tripartite Aβ-Tau-TSPO functional interactions. An astrocytosis also exists in other pathologies, notably in mouse models of multiple sclerosis. However, the knockout of TSPO induces a lower astrocyte reactivity and decreases the severity of symptoms in response to the experimental induction of multiple sclerosis^22^ supporting the pro-inflammatory hypothesis of TSPO. Limitation of astrocyte reactivity by other genetic modifications has also shown ameliorative effects in AD and amyotrophic lateral sclerosis^41, 42, 43^. In AD, an early activation of astrocytes and overexpression of TSPO by astrocytes was observed^16, 17, 18, 19^. All these data converge to support the hypothesis that TSPO in astrocytes could be one of the early determinants of the negative course of AD and, by extension, of some other neurodegenerative pathologies.

The mechanism of action of the absence of TSPO on the decrease in Aβ and Tau does not seem direct since TSPO is predominantly of extra-neuronal origin. However, astrocytes participate in the amplification of the pathology by stimulating the synthesis and spread of Aβ and Tau^44, 45, 46^. This effect could be due to the pro-inflammatory environment created by the stimulation of astrocytes. However, we have shown that the reactivity of astrocytes is not uniformly modified in the hippocampus. Astrocytes of the dorsal part of the hippocampus (dorsal hippocampus including dentate gyrus) are more affected than in the ventral part by the absence of TSPO. Astrocyte reactivity therefore does not appear to be directly related to sAPPα since its expression is stable throughout the dorso-ventral axis of the pyramidal layer and absent in the granular layer of the dentate gyrus. It therefore seems more probable that the presence of Aβ in the parenchyma presents a dorso-ventral gradient which would induce this heterogeneity of the astrocytic response. Supporting this idea, we have shown that the accumulation of Aβ plaques begin in the dorsal hippocampus and subiculum in 3xTgAD mice^29^.

We conducted this study exclusively at the early stages of the disease to determine whether TSPO may represent a treatment target before the onset of the irreversible effects of neuronal death leading to cognitive deficits. Although the absence of TSPO induces beneficial effects, microglia is not affected although it is considerably enriched in genes associated to AD risk-associated genomic loci and may therefore show an early involvement in the pathological cascade of the disease^7^. It is therefore possible that in more advanced stages the absence of TSPO would also induce a modification of microglial response. This difference in temporal course between TPSO from astrocytes and TSPO from microglia has already been observed in various pathologies^47^ and points to the fact that the pathological stage must be considered in the development of therapeutic strategies. Some studies have tested TSPO strategies and seem to show ameliorative effects ^48, 49, 50, 51^ but the presence of off-target effect cannot be excluded ^52, 53^ which can be measured using TSPO^−/−^ mice.

Two limitations must be made. Firstly, the effects we observed at the neuropathological level are not translated by a cognitive improvement, due to the absence of disorders at this age in 3xTgAD mice. An additional study could be conducted with older animals to measure this impact. However, we showed that the absence of TSPO does improve the cognitive effects linked to Tau. Secondly, the use of antibodies for the characterization of Aβ and Tau brings a limit in the interpretation of these data. Indeed, the specificity of antibodies towards the different peptides is still discussed^54^.

Finally, interestingly and advantageously, Aβ42, the most toxic form of Aβ, is the one most affected by the absence of TPSO. Combined with a decrease in Tau pathology and an improvement in one of the neuropsychiatric symptoms, this observation makes TSPO a target with high potential for success as a treatment for Alzheimer’s disease.

## Methods

### Animals and sample collection

Experimental approval was obtained from the ethics committee for animal experimentation of the canton of Geneva, Switzerland. Female WT, TSPO^−/−^, 3xTgAD and 3xTgAD.TSPO^−/−^ mice were reared under standard light/dark conditions with *ad libitum* access to food and water. Animals were randomly assigned and tested blind to both experimental condition and genotype. Under anesthesia, an intracardiac saline perfusion was performed before brain removal. One hemisphere was immersed in formalin for 24 hours and then in a sucrose solution of increasing concentration before being frozen and cut with a cryostat (30 μm sections) for carrying out immunohistochemistry and immunofluorescence. The other hemisphere was dissected to isolate the hippocampus for protein measurement and the cortex for mRNA measurement. The samples were frozen in liquid nitrogen and then stored at −80°C.

### Postmortem human samples

Experimental approval was obtained from the local ethics committee for human experimentation of the canton of Geneva, Switzerland. Human frozen hippocampus come from the Netherland brain bank and formalin fixed paraffin embedded human hippocampus sections (20 μm) come from diagnosed AD subjects of the Geneva Brain Bank^55^. Characteristics of the samples are given in Supp table 1.

### Protein extraction and quantification

Human and mouse samples were sonicated after immersion in a solution of triton (50mM Tris HCl, 150mM NaCl, 1%Triton x100, protease and phosphatase inhibitors 1x, pH=7.4). After centrifugation (20 000 g, 20 min, 4°C), the supernatants collected form the triton soluble fraction. The pellet was taken up with a solution of guanidine (5M guanidine, 50mM Tris HCl, protease and phosphatase inhibitors 1x, pH=8). After gentle agitation (3h, 4°C) and centrifugation (20 000 g, 20 min, 4°C), the supernatants collected form the guanidine soluble fraction. A protein assay was performed by BCA to express the data measured by ELISA as a function of the total amount of proteins and to deposit 20 μg of proteins per sample by western blot.

### ELISA

The determination of the amount in different forms of Aβ40, Aβ42, sAPPα,sAPPβ and Tau was carried out by ELISA from the following kits: Aβ40 human ELISA kit; Aβ42 human ultrasensitive ELISA Kit and Tau (phospho) [pT231] human ELISA Kit from Thermo Fisher, and sAPPα and sAPPβ ELISA Kit from Mybiosource. The procedure was performed according to the instructions of the kits. Briefly, the samples diluted in the dilution buffer were exposed for 2-3 hours to the specific antibody attached to the wells of the 96-well plate. After washing, the primary and then the secondary antibody were introduced into the wells for 30 min before adding the stabilized chromogen (30 min) and the stop solution. The OD reading was performed at 450 nm in the presence of a standard curve, specific to each kit.

### Immunostaining and sholl analysis

Hippocampus sections (4 slices separated by around 200 μm/animal; 4 slices per AD subjects) were incubated with the primary antibody (4°C, overnight), rinsed in 1xPBS, incubated with the secondary antibody (90min, RT), rinsed in 1xPBS and treated with Sudan Black (0.3% in 70% ethanol) before being stained with a DAPI (30 nM, 10min) or Methoxy-XO4 (MxO4) solution (1/500, 30 min, Tocris). To increase the detectability of the AT8 staining, a revelation with 0.2 mg/ml DAB (Sigma-Aldrich) in 1xPBS containing 100 μl/L H2O2 was performed instead of Sudan black and DAPI staining. The procedure was modified to co-stain STAT3 and VIMENTIN as follows: the sections were pretreated with methanol (−20°C, 20 min), rinsed in 1xPBS and then immersed in a blocking solution (1xPBS, 0.3% Triton, 1% BSA) before being shaken for 3 days with the antibody solution (1/200 in Signal Stain). After two series of rinsing in 1xPBS interspersed with exposure to the secondary antibody (1/200 in blocking solution), the sections are mounted using Fluorsave and marked with DAPI (30 nM, 10min). The antibody list was the following: 6E10 (1/200, mouse, Biolegend), A4G8 (1/500, mouse, Biolegend), AT8 (1/1000, mouse, ThermoScientific), GFAP-Cy3 (1/1000, Sigma), HT7 (1/500, mouse, Invitrogen), IBA1 (1/300, Rabbit, Wako), TSPO (1/150, Rabbit Ab100907, Goat Ab118913, Abcam), rabbit anti-mouse HRP (1/100, Dako), goat anti-rabbit Alexa Fluor 488 (1/200, Invitrogen), chicken anti-goat Alexa Fluor 594 (1/200, Invitrogen) and goat anti-mouse Texas Red (1/200, Invitrogen). Images of entire sections were acquired using Zeiss Axioscan for measurement of % of labeled area and fluorescence intensity. The regions of interests have been drawn manually based on anatomy using ImageJ software. Confocal images (2 μm steps) were used for the sholl analysis using the ImageJ software plugging. The soma of astrocytes was manually drawn, and concentric circles were automatically applied (2 μm distance between circles) to measure the number of intersections along the Sholl radii, the area under the curves and the size of soma. The number of animals presenting AT8+ neurons in the hippocampus was manually determined.

### Western blot

Twenty μg of proteins were denatured in 1x Laemmli buffer, 2-5%β-mercapto-ethanol for 10min at 70°C and then migrated with protein ladder (Precision plus protein all blue standards, Bio-Rad) in Bio-Rad gels (Criterion TGX) at 150V for 55min with the manufacturer’s protocol (Bio-Rad). Transfer on LF-PVDF membrane was performed for 7min at 2.5A constant, and up to 25V in the manufacturer buffer using the trans-blot turbo transfer system (Bio-Rad). Membrane was saturated in Tris-buffer (20 mM Tris, 150 mM NaCl, 0.1% tween, pH=7.4) containing 5% non-fat dry milk for 45min and then incubated 48h at 4°C with the primary antibody in Tris-buffer containing 5% non-fat dry milk. Following 3 washes in Tris-buffer (10 min, RT), the membrane was immerged in the secondary antibody in Tris-buffer containing 5% non-fat dry milk for 90 min. Following 3 washes in Tris-buffer (10 min, RT), the fluorescence was detected using the iBright imaging system (ThermoFisher Scientific). The same membrane was reused after a new saturation step. Primary antibodies used at 1/250 were as follows: Actin (Sigma), ApoE (Abcam), BACE1 (Cell signaling), Clusterin (Santa Cruz), GFAP-Cy3 (Sigma), IDE (Abcam), TSPO (Abcam) and secondary antibodies (1/1000) were Alexa Fluor antibodies from Thermo Fisher. Densitometry analysis using ImageJ was performed to quantify proteins and raw data were expressed as function of ACTIN.

### qPCR

Total RNA were extracted using the RNeasy mini kit (Qiagen) and cDNA synthesis were performed using the SuperScript VILO cDNA synthesis kit (Invitrogen) according to manufacturer instructions. Quantitative PCR were performed using Syber green detection and PCR cycles as follows: initial denaturation 95°C, 30sec followed by 40 cycles of 95°C, 15sec; 60°C, 1min and a final step 65°C, 30 sec and then 5 sec to 95°C (0,5°C/sec), 15 sec. Primers used for the detection of mRNA were designed as follows: *App* (fwd:TCAGGGACCAAAACCTGCAT, rev:GCACCAGTTCTGGATGGTCA), *Clu* (fwd: AGCGCACTGGAGCCAAG, rev: TCAGAGACCTCCTGCTCTCC), *Gapdh* (fwd:TGGCAAAGTGGAGATTGTTGCC, rev:AAGATGGTGATGGGCTTCCCG), *Gfap* (fwd:ACGACTATCGCCGCCAACT, rev:GCCGCTCTAGGGACTCGTTC). Relative mRNA expression was normalized to *Gapdh*.

### Behavioral tests

*Elevated plus maze*. Animals were placed at the center of the maze and were free to explore open (20 × 5 cm) and closed (20 × 5 × 30 cm) arms for 5 min. The time past in visiting open arms and the time past performing head-dipping were automatically measured using Ethovision software (Noldus). *Y maze*. Animals were placed in one of the three arms (20 × 10 × 30 cm) and were free to explore for 5 min. The number of good and total alternations was automatically measured using Ethovision software.

### Brain AAV injections

Under isoflurane anesthesia and buprenorphine (Temgesic), a bilateral stereotaxic injection of 2 μl of AAV-Tau or AAV-GFP was performed into the dorsal hippocampus (AP: −3mm, Lat:+/-3mmm, V:-1.5 mm). After injecting at a rate of 0.2 μl/min, the syringe was left in place for 2 min before withdrawal. AAV-Tau was prepared in the laboratory according to the protocol described previously^56^. Briefly, the complete human Tau sequence was cloned and then inserted into pAd / CMV / V5-DEST adenoviral vectors, under the control of a cytomegalovirus promoter (ViraPower Adenovirus Expression System, Invitrogen). The AAV-Tau was verified by sequencing and the production of Tau by western blot. After titration, the viruses were stored at −80°C before their use. Before injection, they were diluted in 0.1 M phosphate buffer saline (PBS) with 0.001% pluronic acid.

### Statistics

A sample size analysis was performed using the graphical Douglas Altman’s nomogram approach with p≤0.05 and β<0.2. Animals were randomized for virus injection. Data production and analysis of raw data of all techniques was done blind to the identification of the samples. Outliers were identified using the ROUT method (maximal false discovery rate = 1%). Comparison between two groups were performed using t-test. The number of mice AT8^+^ was compared using chi-squared. One- and two-way ANOVA were used to analyze the effect of AD stage in human, genotype and age in mice. Data are presented as individual values and mean ± SEM. All statistical experiments were performed with GraphPad Prism 9. The number of stars on graph indicate the level of significance, as follows: *p<0.05, **p<0.01, ***p<0.001, ****p<0.0001.

## Data availability

The datasets generated during and/or analysed during the current study are available from the corresponding author on reasonable request.

## Acknowledgements

We are grateful to Maria Surini and Pia Lovero for the technical assistance. A part of this work was supported by the Velux foundation (grant number 1123).

## Author contributions

KC, TZ, VG, and BBT designed the AAV experiment and KC, ST, PM and BBT designed all the other experiments. KC, LM, FB, QA and BBT performed experiments and analyzed data. R.B.B., G.-J.L. and R.J.M. provided global *Tspo* knockout mice (C57BL/6-Tspo^tm1GuMu(GuwiyansWurra)^) and contributed to the design of experiments. KC, ST, QA, PM and BBT wrote the manuscript. All authors reviewed and edited the manuscript.

## Figures and Figure legends

**Supplemental Fig. 1.**
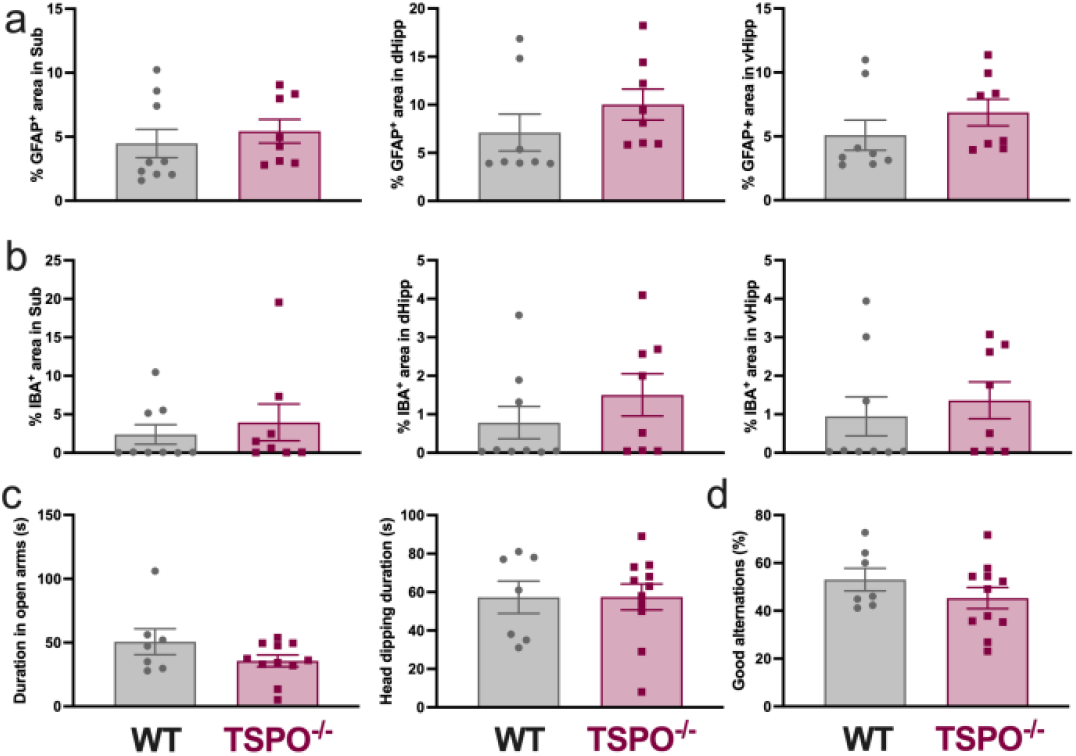
TSPO^−/−^ mice did not show glial activity and behavioral changes. **a**, Quantification of % of positive GFAP-ir area in the subiculum (Sub), dorsal hippocampus (dHipp) and ventral hippocampus (vHipp) (two-tailed unpaired t-test: *P* > 0.05). **b**, Quantification of % of positive IBA1-ir area in Sub, dHipp and vHipp (two-tailed unpaired t-test: *P* > 0.05). **c**, Time past in open arms and head-dipping duration in the EPM (two-tailed unpaired t-test: *P* > 0.05). **d**, % of good alternation in the Y-maze (two-tailed unpaired t-test: p>0.05).

**Supplemental Table. 1.**
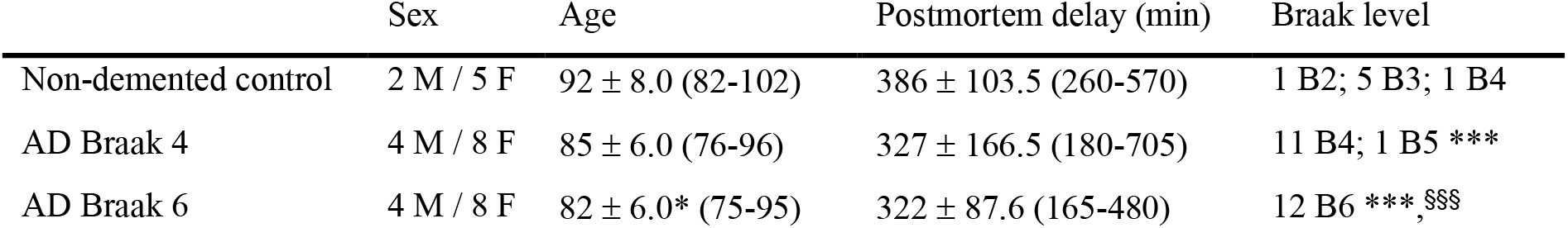
Characteristics of human samples. The three groups did not differ neither in sex distribution (X^2^=0.05, *P* > 0.05) nor in postmortem delay (one-way ANOVA: *F_2,28_* = 0.65, *P* = 0.53). In contrast, age (one-way ANOVA: *F_2,28_* = 4.69, *P* = 0.017, post hoc Holm-Sidak test: AD Braak 4 vs non-demented, *P* = 0.06; AD Braak 6 vs non-demented, \**P* = 0.0156; AD Braak 6 vs AD Braak 4, *P* = 0.38) and Braak level (one-way ANOVA: *F_2,28_* = 214.2, *P* < 0.0001, post hoc Holm-Sidak test: AD Braak 4 vs non-demented, \*\*\**P* < 0.0001; AD Braak 6 vs non-demented, \*\*\**P* < 0.0001; AD Braak 6 vs AD Braak 4, §§§*P* < 0.0001) are different between groups. Age and postmortem delay are presented as mean ± SD and the range. B: braak level; F: females; M: males.

