## Supplementary material for "Knockout of TSPO delays and reduces amyloid, Tau, astrocytosis and behavioral dysfunctions in Alzheimer’s disease": Supp data

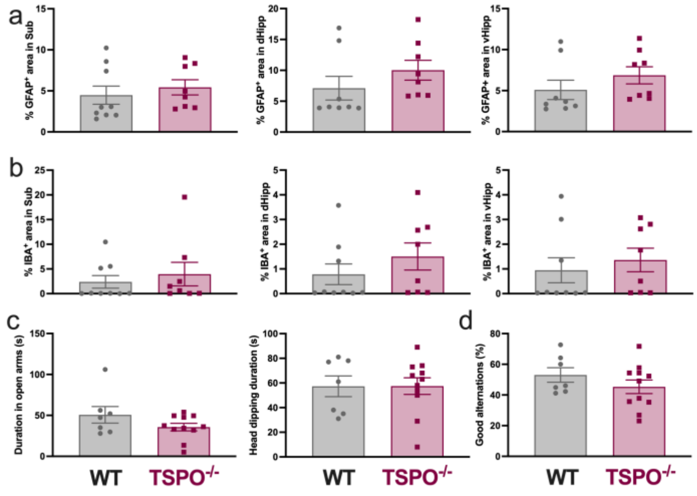


**Supplemental Fig. 1 TSPO$^{-/-}$ mice did not show glial activity and behavioral changes.**

**a,** Quantification of % of positive GFAP-ir area in the subiculum (Sub), dorsal hippocampus (dHipp) and ventral hippocampus (vHipp) (two-tailed unpaired t-test: $P > 0.05$). **b,** Quantification of % of positive IBA1-ir area in Sub, dHipp and vHipp (two-tailed unpaired t-test: $P > 0.05$). **c,** Time past in open arms and head-dipping duration in the EPM (two-tailed unpaired t-test: $P > 0.05$). **d,** % of good alternation in the Y-maze (two-tailed unpaired t-test: p>0.05).

| | Sex | Age | Postmortem delay (min) | Braak level |
|---|---|---|---|---|
| Non-demented control | 2 M / 5 F | 92 ± 8.0 (82-102) | 386 ± 103.5 (260-570) | 1 B2; 5 B3; 1 B4 |
| AD Braak 4 | 4 M / 8 F | 85 ± 6.0 (76-96) | 327 ± 166.5 (180-705) | 11 B4; 1 B5 *** |
| AD Braak 6 | 4 M / 8 F | 82 ± 6.0* (75-95) | 322 ± 87.6 (165-480) | 12 B6 ***,$^{§§§}$ |

**Supplemental Table. 1 Characteristics of human samples.**

The three groups did not differ neither in sex distribution ($X^2$=0.05, $P > 0.05$) nor in postmortem delay (one-way ANOVA: $F_{2,28} = 0.65$, $P = 0.53$). In contrast, age (one-way ANOVA: $F_{2,28} = 4.69$, $P = 0.017$, post hoc Holm-Sidak test: AD Braak 4 vs non-demented, $P = 0.06$; AD Braak 6 vs non-demented, *$P$ = 0.0156; AD Braak 6 vs AD Braak 4, $P = 0.38$) and Braak level (one-way ANOVA: $F_{2,28} = 214.2$, $P < 0.0001$, post hoc Holm-Sidak test: AD Braak 4 vs non-demented, ***$P < 0.0001$; AD Braak 6 vs non-demented, ***$P < 0.0001$; AD Braak 6 vs AD Braak 4, $^{§§§}P < 0.0001$) are different between groups. Age and postmortem delay are presented as mean ± SD and the range. B: braak level; F: females; M: males.
